# Functional inactivation of the telomerase chaperone TCAB1 primes cells for the activation of ALT in osteosarcoma

**DOI:** 10.64898/2026.01.02.691864

**Authors:** Joshua Keegan, Sydney Sorbello, Joakin Mori, Shugo Muratani, Ignaty Leshchiner, Christopher M. Heaphy, Rachel L. Flynn

**Affiliations:** Department of Pharmacology, Physiology & Biophysics Boston, MA 02118, USA; Bioinformatics Program, Faculty of Computing and Data Science, Boston University Boston, MA 02118, USA; Department of Pathology and Laboratory Medicine Boston University School of Medicine Boston, MA 02118, USA; Department of Medicine Boston University School of Medicine Boston, MA 02118, USA

## Abstract

Activation of the alternative lengthening of telomeres (ALT) pathway accounts for cellular immortalization in 75% of pediatric osteosarcoma. ALT does not rely on a single enzyme but instead, catalyzes telomere elongation via homologous recombination. There has been steady progress in defining the mechanisms that regulate the ALT pathway. However, the spectrum of genetic mutations that underlie activation of ALT remains unclear. Osteosarcomas, like many cancers, frequently harbor inactivating mutations in the tumor suppressor gene *TP53*. However, instead of single nucleotide variants that lead to expression of mutant TP53 protein, osteosarcoma tumors often acquire unique structural variations within the first intron of the TP53 gene leading to complete gene inactivation. *TP53* is located on chromosome 17p13.1 in a head-to-head orientation and partially overlapping with the gene *WRAP53* (WD repeat containing antisense to TP53). WRAP53, also known as TCAB1, is an RNA chaperone that is an essential component of the telomerase holoenzyme. TCAB1 functions to facilitate trafficking of the telomerase RNA (hTR) within the nucleus to ensure assembly and localization of the telomerase enzyme to telomere ends to promote telomere elongation. Loss of TCAB1 function abolishes telomerase activity, driving progressive telomere attrition. Here, using whole-genome sequencing of osteosarcoma samples we identified SVs within the *TP53* gene that not only compromise *TP53*, but also inactivate *TCAB1*. These *TCAB1* SVs were prevalent in approximately 40% of ALT positive osteosarcoma tumors suggesting that functional inactivation of the telomerase holoenzyme may be an early and previously unrecognized event contributing to the activation of the ALT pathway.

## INTRODUCTION

Telomeres cap the ends of linear chromosomes and provide a molecular barrier for the human genome. Following each cell division, progressive telomere shortening erodes that barrier and threatens the stability of essential genetic material. Critically short, or dysfunctional telomeres induce replicative senescence and/or cell death and ultimately, lead to cellular aging. To overcome the replicative senescence associated with critically short telomeres, most cancer cells activate one of two known mechanisms of telomere elongation, reactivation of the enzyme telomerase or activation of the alternative lengthening of telomeres (ALT) pathway^1^. A small minority of tumors lack both mechanisms and continue to proliferate even as their telomeres continue to shorten (i.e. ever-shorter telomere phenotype; EST)^2,3^. While the prevalence of EST in human cancer is unclear, reactivation of telomerase accounts for cellular immortalization in approximately 90% of human cancers while the alternative lengthening of telomeres (ALT) pathway is active in approximately 10% of all cancers. However, the prevalence of ALT increases to 80% in tumors of mesenchymal origin, including osteosarcoma^4,5^. Nevertheless, the etiology surrounding the activation of ALT in osteosarcoma remains unclear.

Reactivation of telomerase (hTERT) commonly occurs through promoter mutations, gene amplification, or epigenetic changes and promotes telomere elongation via the action of the telomerase reverse transcriptase enzyme^6–10^. In contrast, the ALT pathway does not rely on a single enzyme, but catalyzes telomere elongation via homologous recombination^11,12^. ALT positive tumors frequently harbor functionally inactivating mutations in genes involved in chromatin remodeling and DNA damage repair, including *ATRX* (α-thalassemia/mental retardation syndrome X-linked), *DAXX* (death-domain associated protein), *SLX4IP* (SLX4 interacting protein), or *SMARCAL1* (SWI/SNF Related, Matrix Associated, Actin Dependent Regulator of Chromatin, Subfamily A-like 1)^13–16^. However, functional inactivation of these proteins alone is insufficient to induce activation of the ALT pathway suggesting that additional defects likely contribute to the process^17–19^. Recent studies have demonstrated that while loss of ATRX in cell culture promotes some phenotypes associated with ALT, combining *ATRX* knockout with knockout of the RNA component of the telomerase catalytic core *hTR,* generates bona fide ALT activity^20^. These data suggest that perhaps functional inactivation of the telomerase holoenzyme may precede activation of ALT in vivo.

The telomerase catalytic core consists of the hTERT reverse transcriptase and the hTR RNA template and together this minimal core complex is sufficient to promote telomerase activity in vitro^21^. However, large scale purification strategies have identified that the telomerase holoenzyme is a much larger ribonucleoprotein complex^22^. Structurally, the hTR RNA consists of a pseudoknot containing the template region bound by hTERT and an H box (consensus ANANNA) and ACA box (H/ACA) motif that acts as a scaffold for recruitment of the small nucleolar ribonucleoproteins (snoRNP) DKC1, NHP2, NOP10, and GAR1. This H/ACA domain also harbors a Cajal body box, or CAB motif that is bound by the protein TCAB1^22,23^. Together, these proteins facilitate the processing and trafficking of the hTR transcript within the nucleus to ensure assembly and localization of an active telomerase enzyme at telomere ends^24^. Loss of function of DKC1, NHP2, NOP10, or TCAB1 leads to defects in hTR maturation and as a result, loss of telomerase activity. Moreover, genetic mutations in DKC1, NHP2, NOP10, and TCAB1 have been identified in patients with the telomere length disorder, dyskeratosis congenita (DC). DC is a rare disease pathology that is defined by a genetic predisposition to critically short telomeres due to deficiencies in telomerase activity^25^. These findings highlight the importance of the telomerase holoenzyme, not just the catalytic core, in the establishment and maintenance of telomere elongation via telomerase.

hTR processing, folding, and localization are essential for the establishment of telomerase activity and ultimately the promotion of telomere elongation^25,26^. TCAB1 facilitates trafficking of hTR within the nucleus to ensure assembly and localization of the telomerase enzyme to telomere ends^27–31^. Notably, TCAB1 is located on chromosome 17p and partially overlaps in a head-to-head orientation with the TP53 tumor suppressor gene. TP53 is one of the most common genetic mutations in human cancer occurring in approximately 42% of tumors^32^. Although the majority of mutations in TP53 lead to loss-of-function the mechanism of inactivation varies across cancer subtypes^33–37^. In pediatric osteosarcoma, TP53 is often inactivated by structural variations (SVs) including focal deletions or translocations at the 17p13.1 locus^35–38^. Here, we have identified SVs that not only compromise TP53, but also inactivate TCAB1, suggesting that pathogenic mutations in the telomerase holoenzyme that compromise function may be an early event in the activation of the ALT pathway in a subset of pediatric osteosarcoma.

## RESULTS

### Loss of TCAB1 expression in ALT positive osteosarcoma

Previous studies have demonstrated an enrichment of breakpoints within the first intron of *TP53* in osteosarcoma^35–38^. In the human genome *TP53* is positioned on the (-) strand located between 7668421-7687490 and *TCAB1* is positioned on the opposing (+) strand between 7686071-7703502 (hg38). This orientation creates a 1,419 nucleotide overlap between *TP53* and *TCAB1*. Structurally, the *TCAB1* gene consists of three alternative promoters annotated within the genome from 5’-3’ as γ, α, and β^39,40^. The γ promoter lies within the first intron of *TP53* and the α promoter overlaps with the first exon of *TP53 (*Figure 1A)^40^. The more distal 3’ beta promoter drives transcription of the mRNA that encodes the TCAB1 protein. Yet, whether the sequence upstream of the β promoter, in the region overlapping with TP53, is critical for TCAB1 expression is unclear.

**Figure 1:**
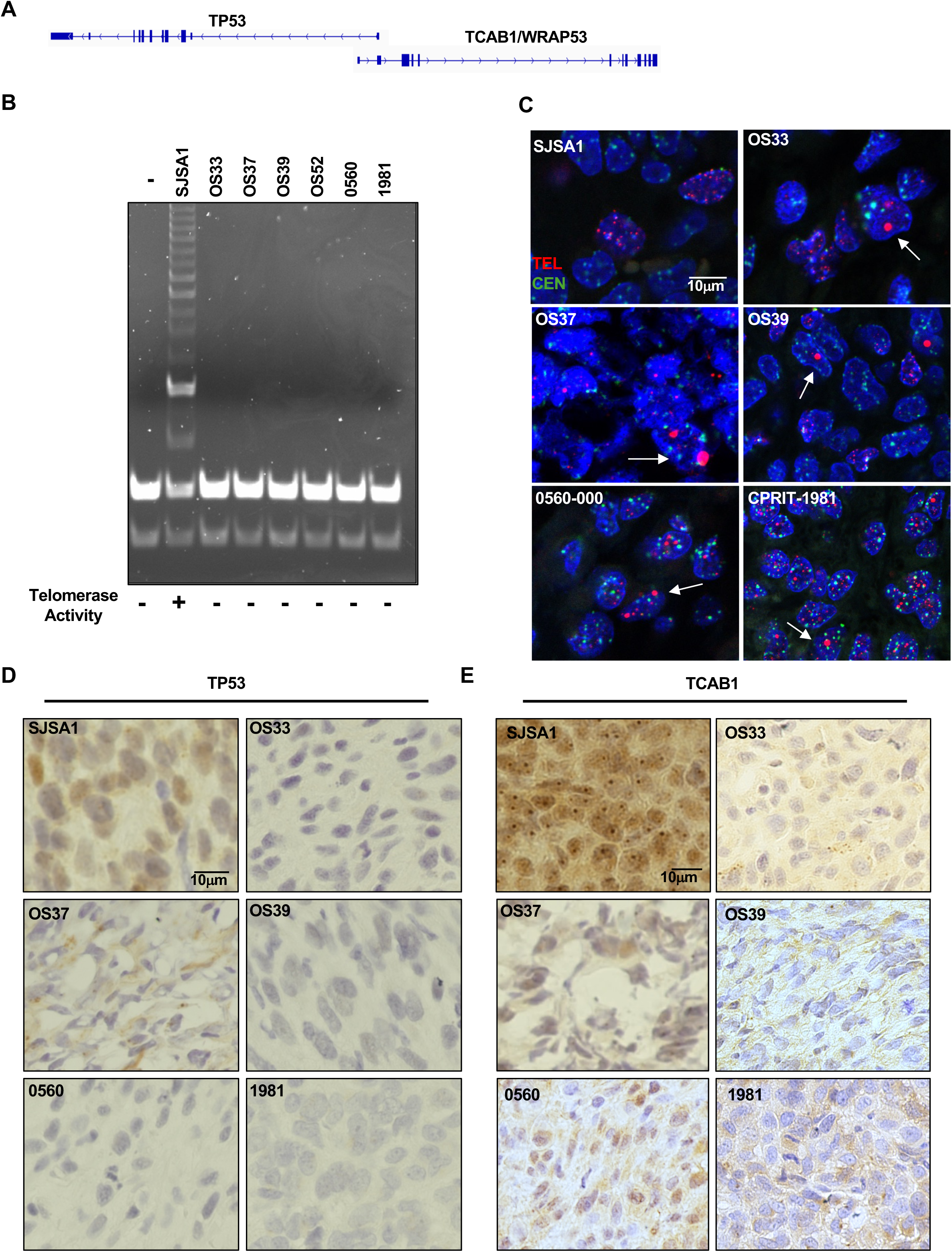
Loss of TCAB1 expression in ALT positive osteosarcoma. A) Schematic representation of the orientation of the *TP53* and *WRAP53/TCAB1* genes within the genome. B) Representative image of an agarose gel from a TRAP assay performed on OS tumors. SJSA1 serves as a telomerase positive control and buffer alone serves as the negative control. IC; internal PCR control, Telomerase positive; +, Telomerase negative; -. C) Representative TEL-FISH result for SJSA1(negative control) and indicated OS tumors. Arrows indicate cells with ultra bright telomeric foci. D) Representative TP53-IHC result for SJSA1 (expresses mutant TP53) and OS tumors with complete loss of TP53 expression. E) TCAB1-IHC result for SJSA1(positive control) and indicated OS tumors with loss of TCAB1 expression.

Given the reported instability within the 17p13.1 locus, we asked whether the observed breakpoints within *TP53* compromise the 1,419 nucleotides overlapping *TCAB1* and ultimately compromise TCAB1 expression. Therefore, we collected a cohort of 16 patient derived xenograft models of osteosarcoma from the Greehey Children’s Cancer Research Institute (GCCRI). We analyzed each tumor for the presence of telomerase activity using a telomere-repeat amplification protocol (TRAP). Remarkably, all 16 tumors analyzed demonstrated complete loss of telomerase activity suggesting they relied on the ALT pathway (Supplemental Figure 1 and Figure 1B). Tumors that rely on ALT are characterized by the presence of ultra bright telomeric foci that can be identified using telomeric fluorescence in-situ hybridization (TEL-FISH). Therefore, to determine ALT status we analyzed formalin-fixed paraffin embedded tumor sections from each sample using the established TEL-FISH assay. Perhaps not surprisingly, all sixteen samples were positive for ultra-bright telomere foci confirming the presence of ALT (Supplemental Figure 1 and Figure 1C).

Previous studies have suggested that approximately 55% of osteosarcoma tumor samples retain SVs within the first intron of *TP53* leading to functional inactivation of the *TP53* gene^36^. To determine whether these SVs also compromise *TCAB1* gene expression, we analyzed both TP53 and TCAB1 protein expression across our cohort of tumors using immunohistochemistry (IHC). Following IHC, we found that 75% of the tumors in our cohort lacked TP53 expression, while 37.5% (6/16) of the cohort demonstrated loss of TCAB1 expression (Supplemental Figure 1 and Figure 1D-E). Moreover, all the tumors demonstrating loss of TCAB1 expression also showed loss of TP53 expression. These data provide early evidence supporting TCAB1 functional inactivation in ALT positive osteosarcoma and suggest that loss of TCAB1 may be a consequence of *TP53* SVs.

### Structural variations in ALT positive osteosarcoma

To further define the structural rearrangements present in our osteosarcoma samples, we isolated genomic DNA from each of the TCAB1 deficient samples (we lacked sufficient material for analysis on OS52) and analyzed each sample using whole genome sequencing (WGS). Using mapped paired-end sequencing reads, we detected structural variants (SV) using three independent SV callers, DELLY, MANTA, and SvABA^41–43^. The results from all three callers were merged using SURVIVOR and annotated using AnnotSV^44,45^. To reduce false positives, we filtered out known common SVs and SVs detected in repeat-rich regions of the genome. The remaining SVs that were identified by ≥2 callers were included in our final dataset. There was an average of 1,464 SVs per sample with deletion events being the most common across all samples (Figure 2A-B).

**Figure 2:**
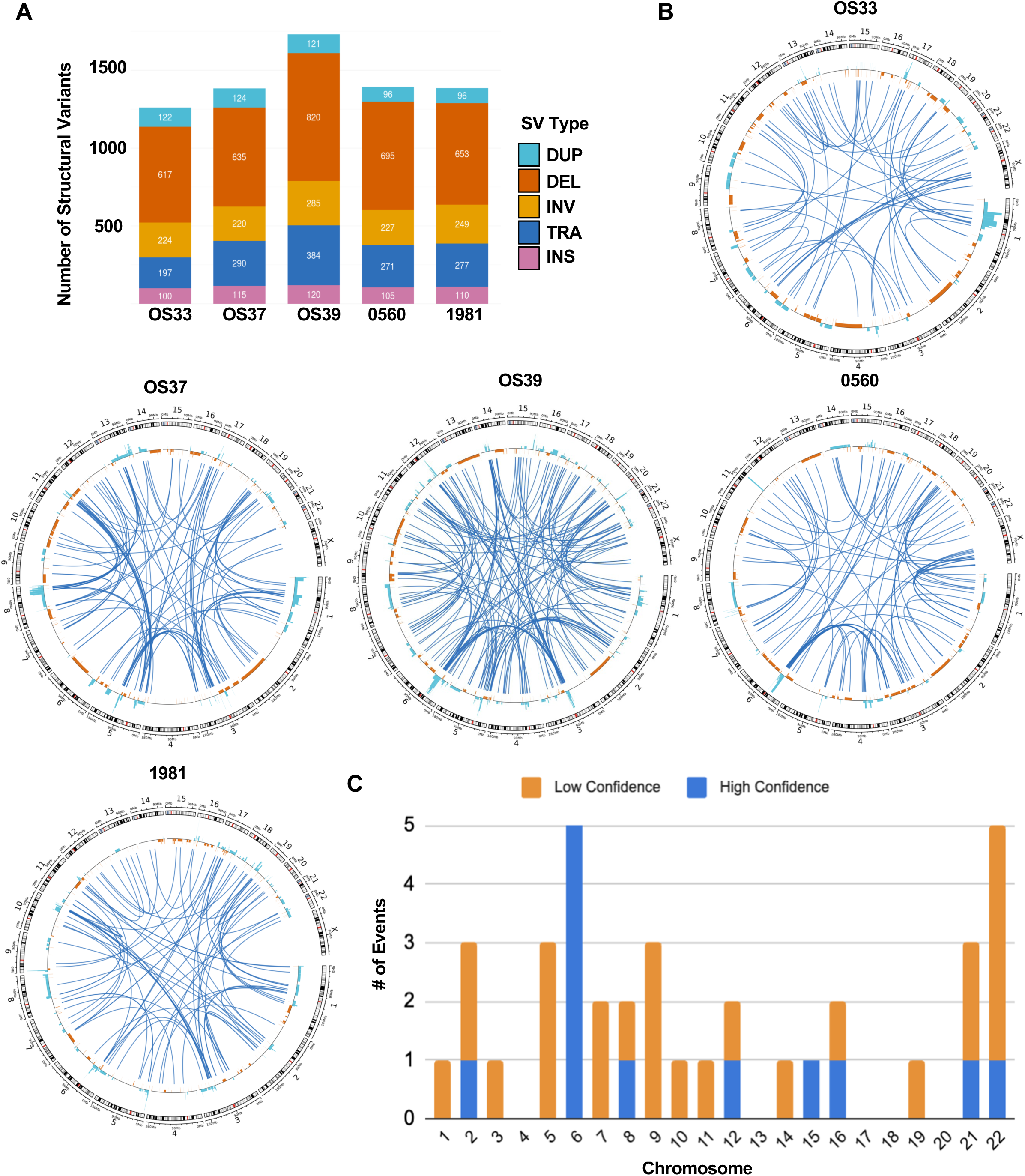
Whole-genome sequencing reveals structural variations in ALT positive osteosarcoma. A) Total number SV per sample represented as a stacked bar plot by the type of SV. Duplication (DUP), deletions (DEL), inversion (INV), translocations (TRA), and insertion (INS) are represented as light blue, orange, yellow, and pink respectively. B) Circos plots for each sample show SV calls across the genome. Duplications, deletions and inversions are represented as tiles in light blue, orange and yellow respectively. Interchromosomal translocations are depicted as dark blue links. C) Frequency of chromothripsis calls across chromosomes for all 5 samples is represented in a stacked bar plot where low confidence and high confidence calls are orange and blue respectively.

In addition to the SVs, we also analyzed somatic copy number alterations using GATK^46^. This analysis revealed copy numbers that deviated from diploid in each sample. Moreover, these copy number alterations demonstrated clear patterns of oscillation often associated with chromosome shattering, or chromothripsis (Figure 2B). To determine whether the instability we observed was a direct result of chromothripsis, we analyzed our WGS data using the computational tool ShatterSeek and chromothripsis events were split into high and low confidence calls. We assigned high confidence calls to chromosomes that had at least 10 total copy-number variations (CNVs) events, 7 or more copy-number oscillation events, and an interchromosomal fragment joining P-value of less than 0.05^47^. We assigned low confidence chromothripsis calls to chromosomes that had at least 10 total CNV events and 4-6 copy-number oscillation events. Overall, the most frequent chromothripsis events were found in chromosome 6 and 22 as we detected a low or high confidence chromothripsis event within these chromosomes in all five samples (Figure 2C). The least frequent chromothripsis events occurred in chromosome 4, 13, 17, 18 and 20 as we did not detect any chromothripsis events in any samples. Thus, while our chromothripsis detection revealed a high level of genomic instability driven by complex rearrangements across samples, chromothripsis alone does not immediately explain the loss of TCAB1 expression on chromosome 17.

### Identification of structural variations in TCAB1 in ALT positive PDX models of osteosarcoma

Although chromosome 17 was not consistently included in chromothripsis events there was clear evidence of rearrangements within the *TP53/TCAB1* locus. In fact, all five samples had evidence of complex SVs in the *TP53* gene and four of five samples had complex SVs in *TCAB1*, with the remaining sample containing a single translocation event within *TCAB1* (Figure 3A). The complex SVs at 17p13.1 spanned both *TP53* and *TCAB1* genes and often consisted of a combination of two or more SVs including inversions, translocations, and deletions in a single gene. As a result, *TP53* and *TCAB1* were often disrupted, not by a single event within the overlapping genomic region, but by multiple SVs along the length of each gene (Figure 3B). In addition to *TP53* and *TCAB1,* we also analyzed genes commonly mutated in ALT including *ATRX, DAXX, SMARCAL1* and *SLX4IP* and found that OS33 contained a complex SV in *ATRX* (Figure 3A).

**Figure 3:**
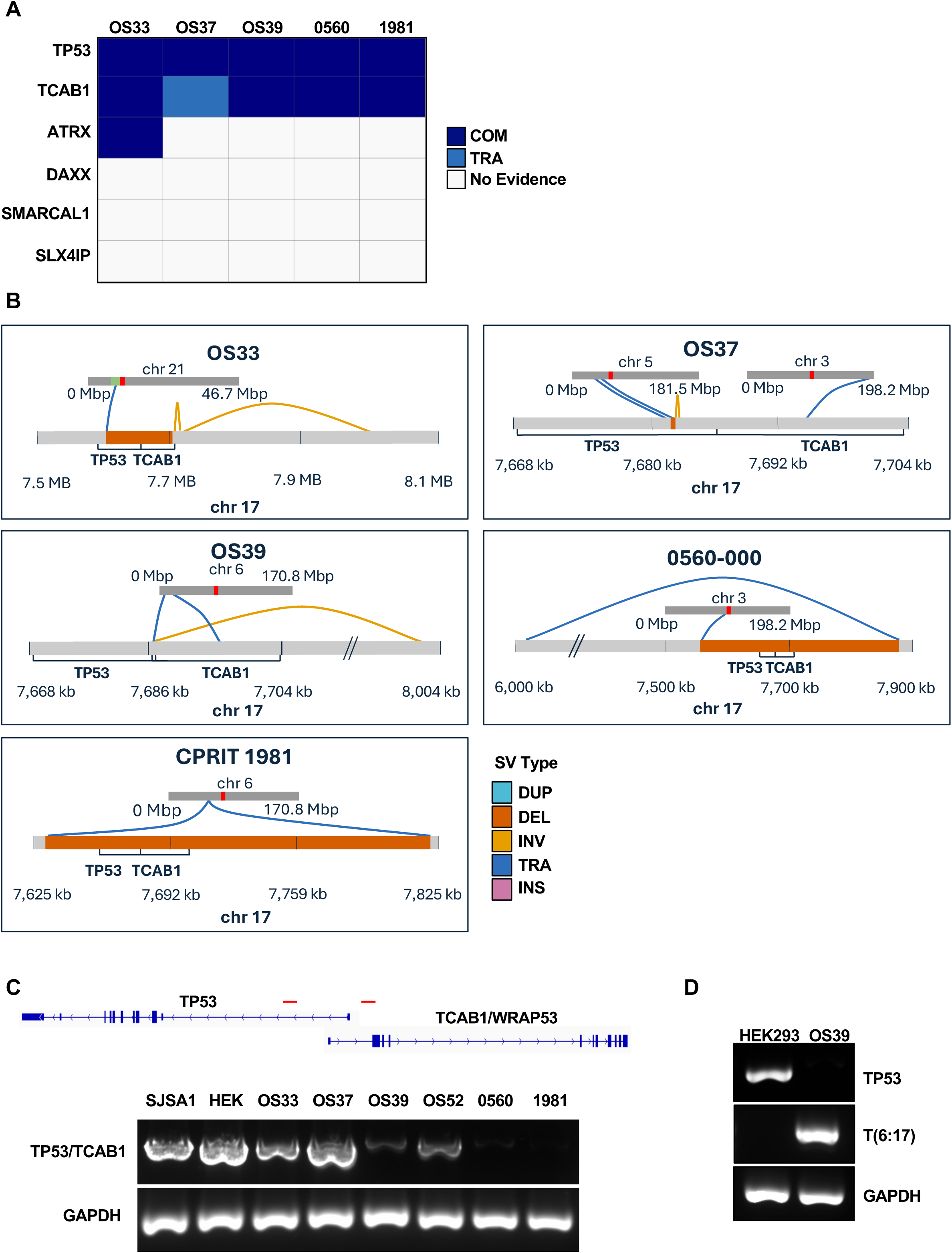
Identification of structural variations in TCAB1 in ALT positive osteosarcoma. A) Categorical heatmap summarizes the SVs for TP53, TCAB1, and ALT associated genes that were identified for each of the five OS tumors analyzed by WGS. Complex (COM) and translocations (TRA) are represented as dark blue and light blue respectively. B) Illustration of the specific SVs identified in each of the five OS tumors analyzed by WGS. C) PCR analysis conducted on the 3.6 KB flanking region between TP53 and TCAB1 that identifies deletion of this locus in tumors 0560-000 and CPRIT 1981. Red lines represent location of primers D) Validation of the translocation event between JARID2-TCAB1 on Ch6:17 in the tumor OS39. HEK293 serves as a negative control for T6:17.

In an effort to confirm the SVs in *TP53* and *TCAB1* identified by WGS, we performed PCR analysis across a 3.6kb region extending from the first intron of *TCAB1* to the first intron of *TP53* in all samples (Figure 3C). As predicted by the SV analysis, we found that the genomic DNA isolated from 0560-000 and CPRIT-1981 tumors failed to produce a PCR product across this 3.6kb region confirming large deletions of the *TP53/TCAB1* locus (Figure 3C). Our initial SV analysis in OS39 predicted a translocation event between the *WRNIP* gene on chromosome 6 and the overlapping region of *TP53* and *TCAB1* on chromosome 17, T(6:17). To confirm this T(6:17) event, we designed primer pairs flanking the break-end and analyzed genomic DNA isolated from either OS39, or the control cell line HEK293, by PCR. As expected, we were unable to detect a PCR product in the genomic DNA isolated from the control HEK293, but we were able to confirm a single band of the predicted size in the genomic DNA from OS39 confirming the T(6:17) translocation (Figure 3D). Genomic DNA isolated from OS33, and OS37 generated the 3.6kb PCR product, but SV analysis clearly defines complex rearrangements in this region. Finally, we were unable to analyze OS52 by WGS, but there was clear loss of TCAB1 protein expression by IHC and poor amplification of this region by PCR confirming functional inactivation of *TP53/TCAB1*.

### Identification of structural variations in TCAB1 in primary osteosarcoma tumors from the TARGET-OS Project

To further assess the prevalence of *TCAB1* mutations in a larger cohort of osteosarcoma samples, we analyzed publicly available WGS data generated by the Therapeutically Applicable Research to Generate Effective Treatments (TARGET) project. Within this project there were 23 additional osteosarcoma patient cases and 21 included quality WGS data. Therefore, we used our computational pipeline to identify both SVs and SNVs within *TP53* and *TCAB1*, and the ALT-associated genes *ATRX, DAXX, SMARCAL1,* and *SLIX4IP*. As expected, we identified ALT positive samples with mutations in *ATRX* and *SMARCAL1*. The mutation in *ATRX* was a deletion whereas the mutation in SMARCAL1 was a SNV generating a premature stop codon (Q653X) (Figure 4A)^48^. Recent studies used a computational tool TelFusDetector to predict ALT status in this cohort of tumors^38,49^. While this prediction method is different than our tissue-based ALT assay, it nevertheless allowed us to compare SVs in our genes of interest between ALT positive (n=10) and ALT negative (n=11) samples. Here, we demonstrate that approximately 50% of the ALT positive samples and 36% of the ALT negative samples showed SVs in *TP53*. Our initial analysis also identified several samples with complex SVs in *TCAB1* that were predicted to lead to functional inactivation. However, there were 3 samples with a single SV event in TCAB1 predicted to disrupt transcription of the α (ENST00000431639.6) and γ (ENST00000457584.6) transcript variants, but not the β variant (ENST00000396463.7) suggesting that these SVs may not affect TCAB1 expression. In fact, analysis of RNA sequencing data that corresponded to 2 of the three samples (sample TARGET-40-0A4I9I did not have corresponding RNA-seq data) confirmed that TCAB1 RNA expression levels were similar to the matched normal controls (Supplemental Figure 3). Together, our data demonstrate that approximately 35% of the ALT positive samples and none of the ALT negative samples, harbor SVs in *TCAB1* (Figure 4B) highlighting SVs in *TCAB1* as a previously unrecognized genetic mutation in ALT positive osteosarcoma (Figure 4C).

**Figure 4:**
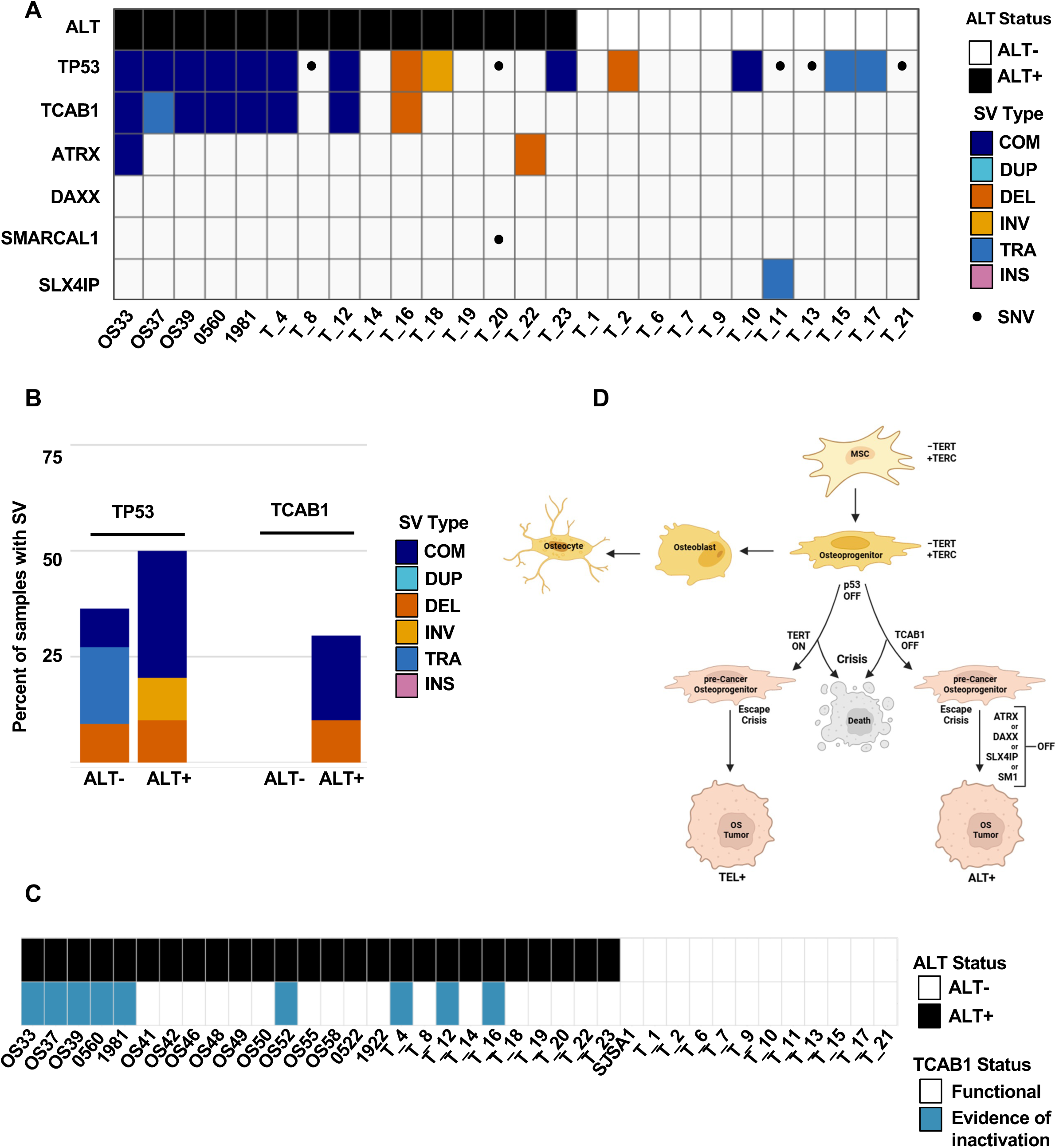
Functional Inactivation of TCAB1 in ALT positive osteosarcoma. A) Categorical heatmap summarizes ALT status, SVs for TP53, TCAB1, and ALT associated genes that were identified for all OS tumors analyzed by WGS. B) Relative percentage of ALT- and ALT+ samples with a SV affecting the TP53 and TCAB1 genomic regions are represented as a stacked bar chart where deletions, inversions, translocations and complex events are orange, yellow, light blue and dark blue respectively. C) Categorical heatmap summarizes the functional inactivation of TCAB1 and ALT status across all 38 samples analyzed by either IHC or WGS. D) Theoretical framework for the evolution of telomerase and ALT positive tumors during osteosarcomagenesis.

## DISCUSSION

In this study we analyzed two independent cohorts of osteosarcoma tumors (GCCRI and TARGET-OS) for SVs within the *TP53/TCAB1* locus. Our findings not only expand on the current literature demonstrating frequent SVs in TP53 in osteosarcoma, but also now highlight SVs in the *TCAB1* gene. These data suggest that *TCAB1* SVs may be an early event in tumor progression coinciding with *TP53* inactivation. Continued cellular proliferation in the presence of critically short telomeres drives TP53 deficient cells into crisis^50^. Although most cells undergo cell death in crisis, rare cells can reactivate telomerase, promote telomere elongation, and emerge from crisis as immortal clones^51^. However, tumors with SVs extending into *TCAB1* cannot support telomerase activity and thus, may undergo selective pressure early on in tumor evolution to activate ALT. Therefore, we propose a shift in the paradigm for the evolution of ALT in a subset of osteosarcoma where genomically unstable osteoprogenitor cells face a ‘fork-in-the-road’ during transformation that hinges on the gain, or loss, of telomerase activity (Figure 4D). Tumors that reactivate hTERT become telomerase positive osteosarcoma whereas tumors that gain complex SV in *TP53/TCAB1* immediately lack any telomere length maintenance mechanism. TCAB1 deficient tumors that acquire additional pathogenic mutations (i.e. *ATRX, DAXX, SMARCAL1, SLX4IP*) may ultimately activate ALT whereas, TCAB1 deficient tumors that do not gain these mutations could either progress to an EST phenotype or simply undergo cell death. In fact, there were two ALT negative tumors with SVs in *TCAB1*. Although the original tumors from the TARGET-OS project are not available for additional analysis, identifying these types of tumors in future cohorts will allow us to match the genetic changes with the acquired cellular phenotypes to further support our understanding of telomere length maintenance mechanisms in osteosarcoma.

Given that mutations in *TCAB1* have not yet been described in cancer genomes, we extended our analysis to the TCGA and found that the TCGA Sarcoma cohort had the highest frequency of *TCAB1* mutations than any other cancer. Interestingly, this cohort does not include osteosarcoma, but instead consists largely of leiomyosarcoma (39.2%), dedifferentiated liposarcoma (23.1%), undifferentiated pleomorphic sarcoma (19.6%), and myxofibrosarcoma (9.8%). These tumors are often diagnosed in adults, but like pediatric osteosarcoma, the prevalence of ALT is high (leiomyosarcoma ∼60% ALT+, dedifferentiated liposarcoma ∼30% ALT+, undifferentiated pleomorphic sarcoma ∼65%, and myxofibrosarcoma ∼75%). Using this, we compared the copy number variations (CNVs) at 17p13.1 in the sarcoma cohort to CNVs at 17p13.1 in the breast cancer cohort where the prevalence of ALT is less than 4%. Here, we found a clear and unique pattern of focal deletions in sarcomas that were not present in breast cancer (Supplemental Figure 4). Although we cannot stratify this cohort based on ALT status, the data demonstrate unique alterations within the *TP53/TCAB1* locus in sarcomas that are not often observed in other tumor types, suggesting that the cellular context likely contributes to the unique mechanism of TP53/TCAB1 inactivation and activation of ALT in these tumor types.

Osteosarcoma is a rare primary bone cancer with an incidence of about 0.48 per 100,000^52^. The 5-year relative rate of survival for patients across all stages of diagnosis is 59%^52–54^. Outcomes have remained stagnant over the past forty years and there have been no new drugs developed in the past 10 years^55–57^. These data highlight an urgent need to further define the complex biology of osteosarcoma and identify tractable targets for therapeutic development. While complex SVs in *TP53* and ultimately *TCAB1*, don’t completely account for the activation of ALT in all osteosarcoma tumors the instability on chromosome 17 highlights previously uncharacterized selective pressures that may arise in the cell providing an opportunity to reexamine ALT dependencies. Our data raise several outstanding biological questions related to the cause of the genome instability, the unique context of the genomic instability, and the contribution of this instability to tumor cell fate. Conceivably, defining the causes and/or consequence of this instability may allow us to identify genetic vulnerabilities in these tumor cells that provide an opportunity to develop targeted therapies for the treatment of osteosarcoma.

## ACKNOWLEDGMENTS

We are grateful to the Greehey Children’s Cancer Research Institute (GCCRI) Patient-Derived Xenograft (PDX) Core and Technical Director Anna Rogojina for providing PDX tissue samples. We thank members of the Flynn and Ganem labs for helpful discussion and the BUMC Cellular Imaging Core facility including Technical Director, Dr. Michael T. Kirber. This publication was supported by the National Center for Advancing Translational Sciences, National Institutes of Health, through BU-CTSI Grant Number 1UL1TR001430, and the Genome Science Institute at Boston University. Its contents are solely the responsibility of the authors and do not necessarily represent the official views of the NIH. R.L.F was supported by R01CA201446, an Edward Mallinckrodt Junior Foundation Award, and a Peter Paul Professorship. C.H. and R.L.F were supported by the US Department of Defense Rare Cancer Research Program grant HT9425-23-1-0819. C.H. was supported by the American Cancer Society (Boston University-Boston Medical Center Pilot and Feasibility Program).The funders had no role in study design, data collection and analysis, decision to publish, or preparation of the manuscript.

## AUTHOR CONTRIBUTIONS

Conceptualization, JWK, SS, RLF, CH, and IL; Investigation Methodology, JWK, SS, SM, JM; Data curation, SS and IL; Formal Analysis, JWK, SS, SM, JM, and IL; Validation, JWK, SS, RLF, IL, CH; Funding acquisition, RLF and CH; Software Supervision, RLF; Project Administration, RLF, CH and IL. Visualization JWK, SS, and RLF; Writing JWK, SS, RLF, CH, JM, SM, and IL.

## MATERIALS AND METHODS

### Patient Derived Xenografts

Early passage, viably frozen PDX tumor samples and FFPE sections were obtained courtesy of Dr. Peter Houghton and and Technical Director Dr. Anna Rogojina at the Greehey Children’s Cancer Research Institute (GCCRI). The FFPE sections for each tumor were provided by GCCRI, except for sample OS37. A frozen tissue block was obtained for OS37, embedded in OCT, and 5μm frozen sections were created. Frozen sections were air dried and then fixed for 10 min in 4% PFA. All staining protocols were followed similarly as with FFPE sections, although deparaffinization and antigen retrieval steps were omitted.

### Telomere-repeat amplification protocol (TRAP)

TRAP assays were performed using the TRAPeze telomerase detection kit (Millipore) according to the manufacturers’ recommendations. PDX samples (25mg) were resuspended in 1x CHAPS Lysis buffer and incubated on ice for 30 min. Lysates were centrifuged at 12,000x g for 20 min at 4°C and protein concentration was determined using Bradford reagent. Approximately, 200 ng of total protein was used in each extension reaction. Internal controls and extension products were amplified by PCR as recommended by the manufacturer. DNA products were separated by 10% PAGE in 0.5x TBE run at 200 V for 2 hr, counterstained with GelRed (1:3,333 for 30 min at room temperature), and visualized with the BioRad ChemiDoc XRS+.

### Assessment of Alternative Lengthening of Telomeres (ALT)

ALT status was assessed using a telomere-specific fluorescence in situ hybridization (FISH) assay on formalin-fixed, paraffin-embedded tissue sections, as previously described^58,59^. Briefly, 5 µm thick slide sections were incubated for 10 min at 65 °C, deparaffinized three times for 5 min in three different xylene containers, and then sequentially rehydrated in 100%, 95%, 70% ethanol, and double-distilled H₂O. Slides were then washed briefly in 1% Tween-20 before being steamed in pH 6.0 citrate-based unmasking solution (Cat# H-3300, Vector Laboratories) for 30 min. After rinsing in double-distilled H₂O, slides were dehydrated through successive ethanol washes (70%, 95%, and 100%). For telomere detection, slides were incubated at 84 °C for 5 min with a Cy3-labeled peptide nucleic acid (PNA) probe complementary to the mammalian telomeric repeat sequence ([N-terminus to C-terminus] CCCTAACCCTAACCCTAA; Cat# F1002, Panagene) and hybridized overnight in a dark, humidified chamber. To verify hybridization efficiency, an Alexa Fluor 488-labeled PNA probe targeting centromeric DNA repeats (Cat# F3012, Panagene) was included in the hybridization mixture. Nuclei were counterstained with DAPI (4′,6-diamidino-2-phenylindole; Sigma-Aldrich) and coverslips mounted using ProLong™ antifade mounting medium (Invitrogen). Slides were analyzed using a fluorescence microscope. Cases were classified as ALT-positive by three independent evaluators based on established criteria, defined by the presence of large, ultrabright intranuclear telomeric foci in ≥ 1% of tumor cells. Areas of necrosis were excluded from evaluation^5^.

### Immunohistochemistry

Five micrometers (5 µm) thick formalin-fixed, paraffin-embedded tissue section slides were deparaffinized using xylene and hydrated through a grade ethanol series prior to antigen retrieval. TCAB1/WRAP53 and p53 immunohistochemistry were performed manually. For TCAB1/WRAP53, slides were steamed in pH9.0 Tris EDTA unmasking solution (Cat# 10-0046, Genemed) for 30 min, blocked with ReadyProbes Endogenous HRP and AP Blocking Solution (Invitrogen, #R37629) and Protein Block (#ab64226, abcam), then were incubated overnight at 4C with rabbit polyclonal primary antibody against TCAB1/WRAP53 (1:100 dilution, Cat# 14761-1-AP, Proteintech). Signals were detected using HRP-conjugated Anti-Rabbit IgG (Cat# PV6119, Leica). For p53, slides were steamed in pH6.0 citrate-based unmasking solution (Cat# H-3300, Vector Laboratories) for 30 min, blocked with ReadyProbes Endogenous HRP and AP Blocking Solution (Invitrogen, #R37629) and Protein Block (#ab64226, abcam), then were incubated overnight at 4C in mouse monoclonal primary antibody against p53 (1:50 dilution, Cat# sc-126, Santa Cruz Biotechnology). Signals were detected using HRP-conjugated Anti-Mouse IgG (Cat# PV6114, Leica). For both TCAB1 and p53, signals were visualized by 3,3′ diamino-benzidine (Cat# D4293, Sigma) prior to Mayer’s hematoxylin counterstaining (Cat# TA-125-MH, Thermo Scientific), dehydration, and mounting. For scoring, retained protein expression was defined as nuclear staining within the tumor cells, and loss of protein expression was defined as the lack of immunolabeling in the tumor cells.

### Primers

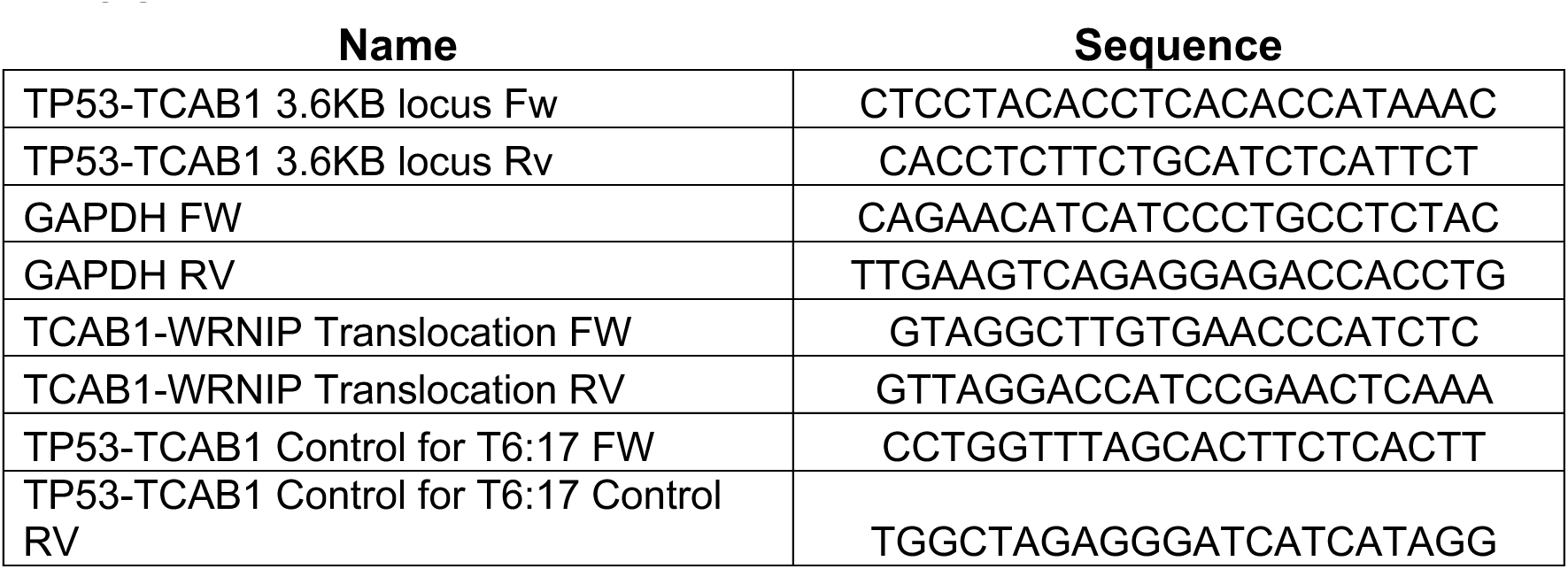

### Polymerase chain reaction (PCR)

PCR was performed on 200 ng of gDNA using the Phusion High-Fidelity PCR Master Mix with GC Buffer (Fisher). The TP53-TCAB1 locus was amplified with the following protocol: 98C for 30 sec, 98C for 10 sec, 62.1C for 30 sec, 72C for 210 sec (repeat steps #2-4 a total of 35 times), and 72C for 10 min. The amplified DNA products were separated by 1% agarose containing GelRed, in 1X TBE, and run for 45min at 100V.

### Genomic DNA (gDNA) Isolation from GCCRI PDX

gDNA from PDX samples (25mg) were isolated according to the QIAamp DNA Mini Kit protocol.

### TARGET-OS

The results published here are in whole or part based upon data generated by the Therapeutically Applicable Research to Generate Effective Treatments (TARGET) (https://www.cancer.gov/ccg/research/genome-sequencing/target) initiative, phs000218. The data used for this analysis are available at the Genomic Data Commons (https://portal.gdc.cancer.gov). dbGaP accession phs000468.v18.p7

21 samples from the TARGET Osteosarcoma (TARGET-OS) cohort were sampled based on their availability of WGS data. These samples were downloaded from the GDC repository using the GDC-client tool (v2.3) [Heath and Ferretti, 2021].

### WGS Data Processing

CRAM files were converted to paired-end FASTQ files. Xengsort (v2.0.8) was used to remove mouse-derived reads from the sequencing data^62^. SeqKit (v2.10.0)^63^ was used to assess read counts and BBMap (v39.26)^64^ was used to correct paired-end FASTQ files that became disordered or unpaired during xengsort classification. Trimmomatic (v0.39)^65^ was used to remove residual unpaired reads. BWA (v0.7.19)^66^ was used to align paired-end FASTQ files to the human reference genome (hg38) with output as BAM file format. SAMtools (v1.21)^67^ was used to evaluate read mapping quality, sort reads by name and coordinate, fix mate reads and mark duplicate reads. The WGS sequencing data for OS33, OS37, OS39, CPRIT-1981, and 0560-000 have been deposited in NCBI’s Sequence Read Archive (SRA) BioProject accession PRJNA1370524. The code developed to analyze PDX WGS data is available at https://github.com/BU-BMSIP/Flynn_WGS_Analysis.

### Structural Variant Detection

A BAM file was used as input for three structural variant (SV) detection tools: Manta (v1.6.0)^43^, Delly (v1.3.3)^42^, and SvABA (v1.2.0)^41^. The results were merged using SURVIVOR (v1.0.7)^45^ retaining SVs detected by at least two of the three tools and allowing a maximum positional distance of 50 bp between matching calls. The SV calls were further subject to filtering parameters by removing common SVs according to the gnomAD database (v4.1)^68^, calls within repetitive regions using a bed file from RepeatMasker (v4.0.6)^69^. Finally, BCFtools was used to remove SVs outside of the canonical chromosomes or with SV lengths greater than 1Mb. Final SV calls were annotated for pathogenicity using AnnotSV (v3.4.6)^44^.

### Copy Number Variant Detection

Copy Number Variant analysis was performed using GATK’s CNV somatic workflow (v4.6.2.0)^46^. In order to perform CNV analysis, 5 normal samples were collected from NCI’s Genomic Data Commons (GDC) (v2.3). The samples were selected from the TARGET ALL Phase 2 cohort. A Panel of Normals (PoN) was constructed from the 5 matched normal samples using GATK CNV somatic panel workflow. Copy Number Variant analysis was performed using GATK’s CNV somatic pair workflow. The final model segment calculations from GATK CNA were used as input for the R package DoAbsolute (v2.2)^70^.

### Chromothripsis Analysis

Chromothripsis analysis was performed using ShatterSeek (v1.1)^47^. The final vcf file was parsed in R in order to obtain necessary information for ShatterSeek and the output tsv from ABSOLUTE was parsed to provide CNV data. High and low confidence chromothripsis calls were determined manually by inspecting the table output from ShatterSeek. High confidence calls were assigned to chromosomes that had at least 10 total CNV events, 7 or more CN oscillation events, and an interchromosomal fragment joining p-value of less than 0.05. Low confidence chromothripsis calls were assigned to chromosomes that had at least 10 total CNV events and 4-6 CN oscillation events.

### Single Nucleotide Variant Analysis

Single nucleotide variants (SNVs) were detected using GATK Mutect2 (v4.6.0.0)^46^. SNV calls were kept if they had 15 or more supporting reads, median mapping quality at least 40 and a filter with value “PASS”. GATK VariantAnnotation was used to annotate the SNVs for their dbSNP ID and ANNOVAR (v2025Mar02)^71^ was used to annotate SNV calls for

### RNA-seq Analysis of TARGET-OS Cohort

The RNA-seq counts matrices and BAM files corresponding to the TARGET-OS samples used in the WGS analysis were downloaded via GDC-client (except sample TARGET-40-0A4I9I which was unavailable)^60^. Differential expression analysis was performed comparing ALT- and ALT+ samples using DESeq2^61^. Results were verified via visual inspection of the BAM files.

---

**Supplemental Figure 1:**
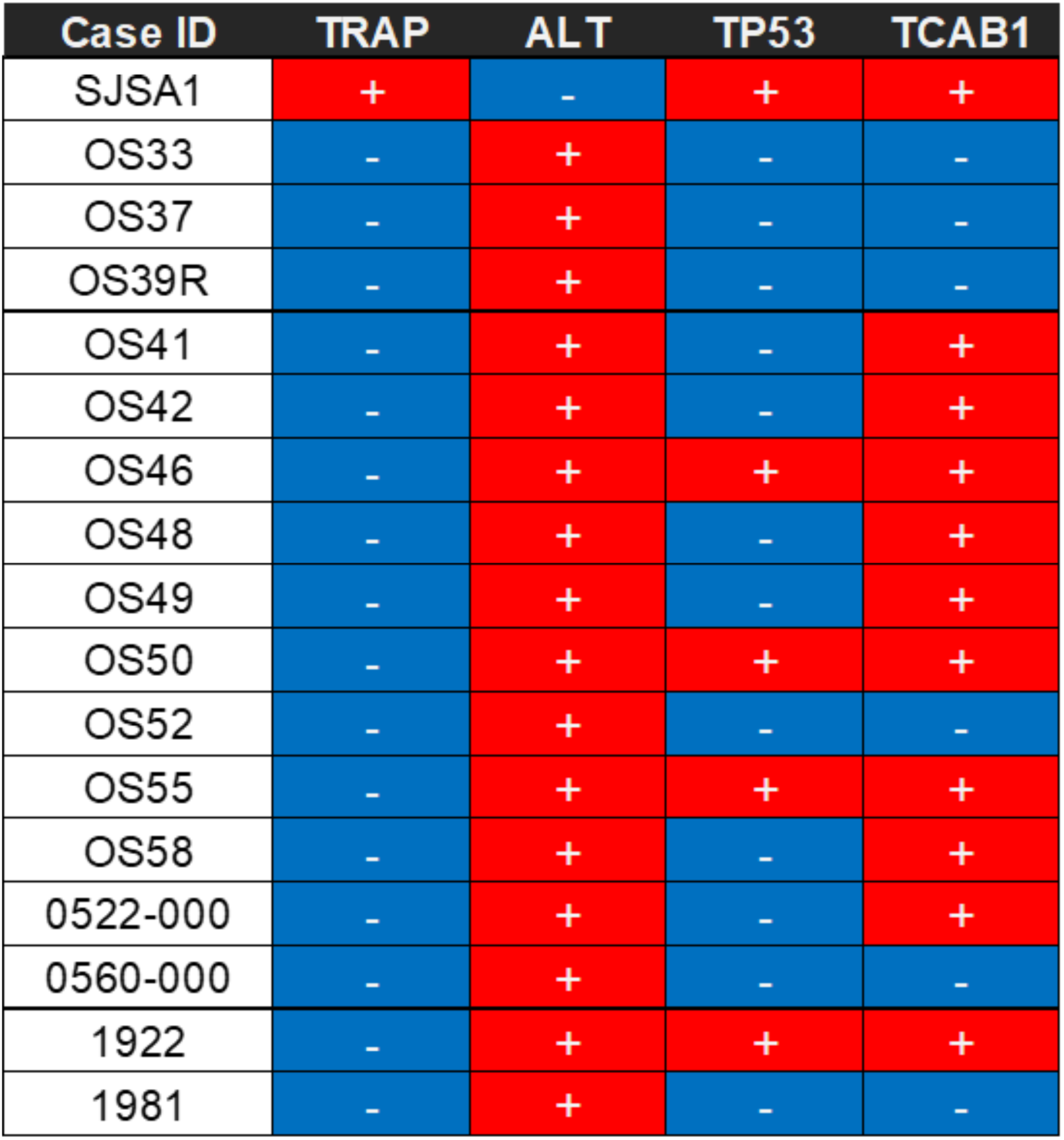
Table summarizing TP53, TCAB1, TRAP, and ALT analysis for the **16 PDX osteosarcoma models from Greehey Children’s Cancer Research Institute (GCCRI).** Columns for TP53 and TCAB1 correspond with IHC analysis, TRAP corresponds to telomerase activity, and ALT corresponds to TEL-FISH analysis. Red, positive; Blue, negative.

**Supplemental Figure 2:**
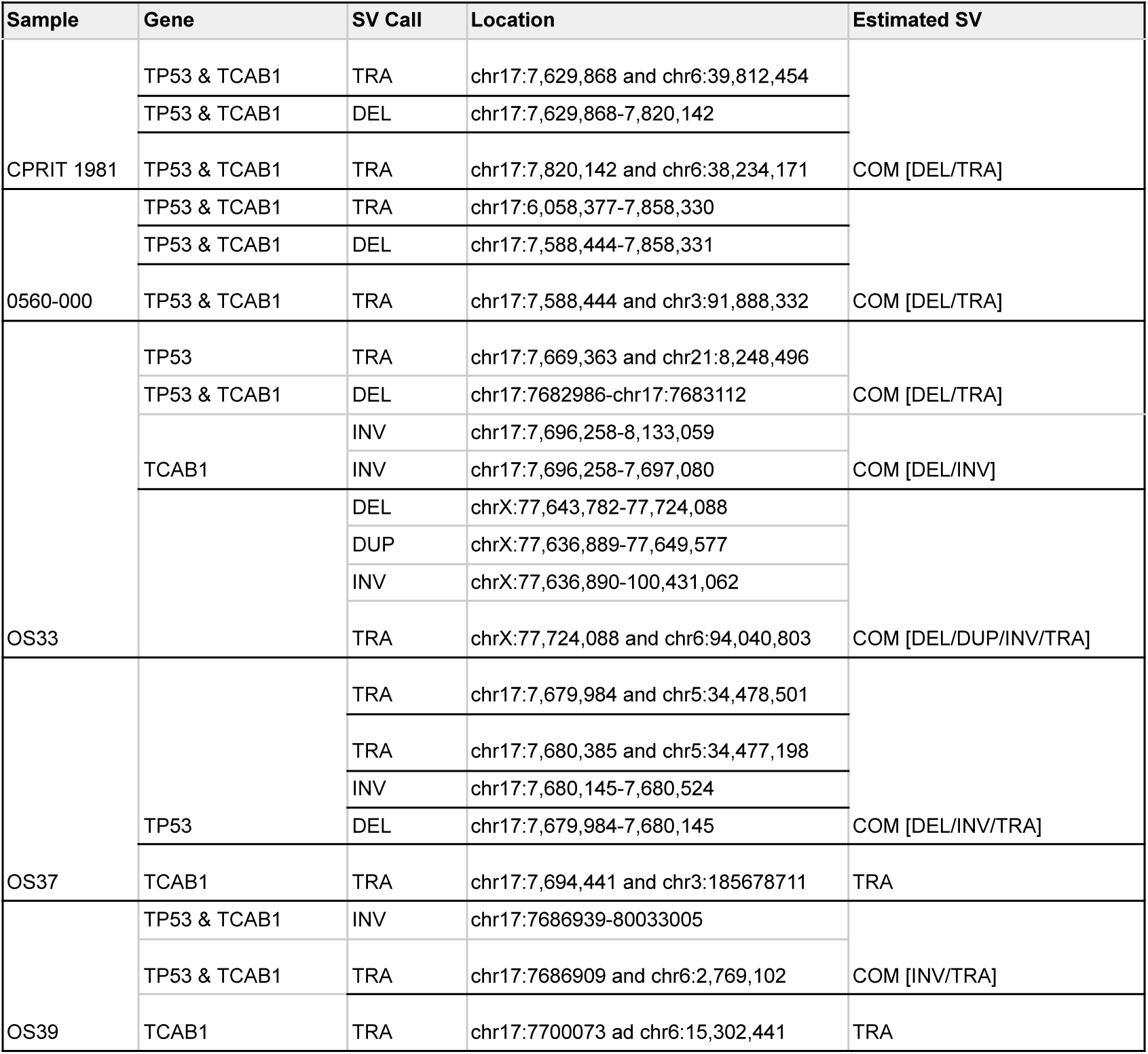
Table summarizing SV in the 5 ALT positive PDX osteosarcoma models analyzed by WGS. The gene, SV call, location within hg38, and overall summary of SV is indicated for each sample.

**Supplemental Figure 3:**
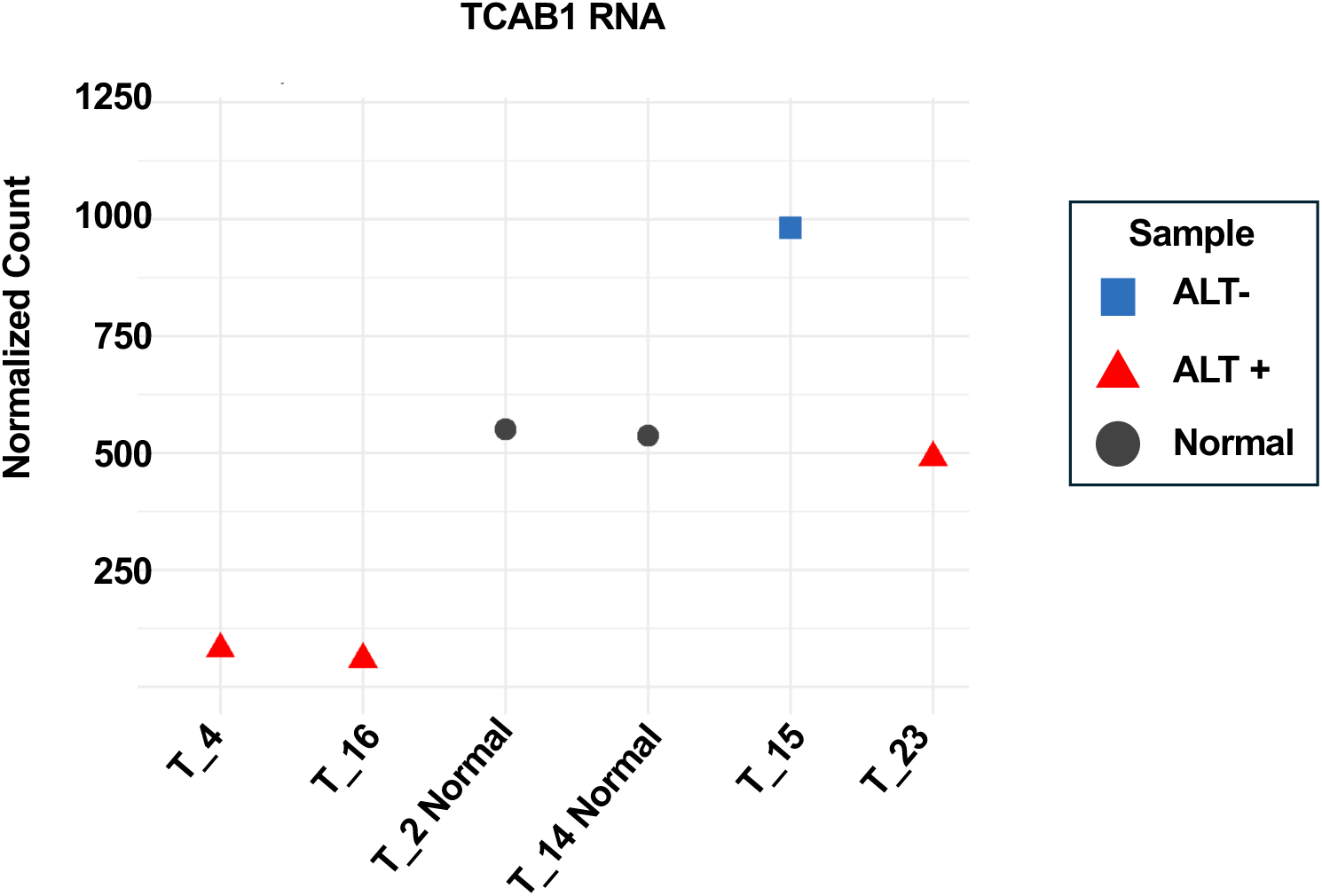
Relative TCAB1 expression in osteosarcoma samples with upstream SVs. Relative expression of TCAB1 in T_4, T_16, T_2 Normal, T_14 Normal, T_15, and T_23 samples following RNA sequencing.

**Supplemental Figure 4:**
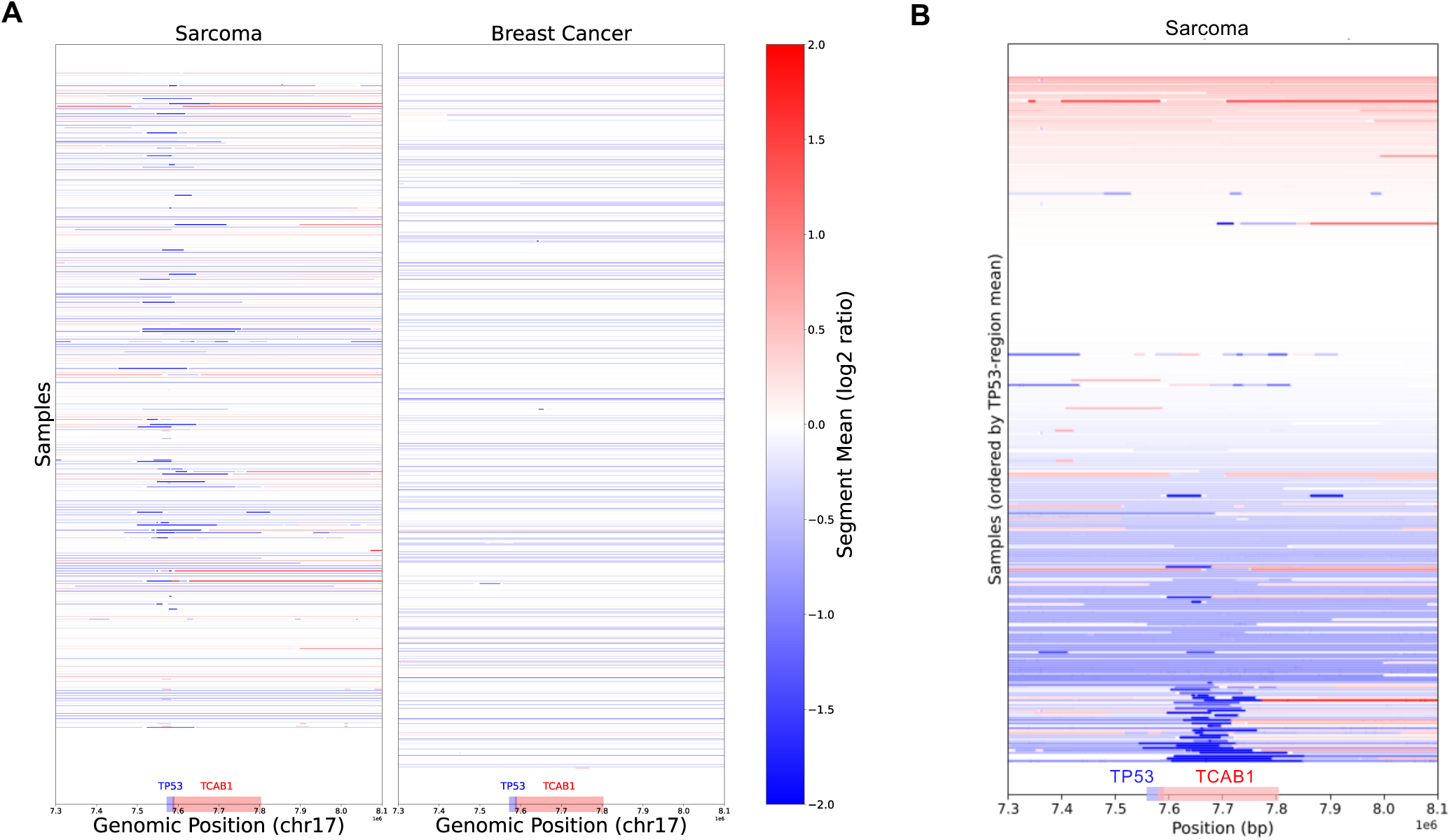
TP53/TCAB1 copy number variations in TCGA cohorts. A) copy number landscape at TP53/TCAB1 locus in TCGA Sarcoma and Breast Cancer cohorts. B) Copy-number landscape for the TCGA sarcoma cohort (n = 272 samples), ordered by mean allelic copy-number at the TP53/TCAB1 locus. CNV segments on chromosome 17 (7.30–8.10 Mb). Arrows indicate samples that also have mutations in ATRX.

